# Leveraging PDE solver for predicting transient space-use dynamics in ecological and epidemiological systems

**DOI:** 10.1101/2022.10.26.513924

**Authors:** Yun Tao

## Abstract

Model predictions of animal and human space-use patterns stemming from individual-level movement behaviors have not only contributed significantly to our understanding of population and community dynamics, but they could also inform the development of conservation, natural resource management, and disease control policies. The recent proliferation of high-resolution movement data has ushered in a paradigm shift in how space use is considered: instead of being defined narrowly as the stationary, long-term distribution of individual locations, there is a growing recognition of its transient dynamics, e.g.., how space-use pattern varies before it eventually stabilizes. However, movement models are slow to follow due to longstanding technical challenges in solving transient space-use dynamics. Here, we introduce a numerical framework that enables transient analysis of mechanistic movement models based on partial differential (Fokker-Planck) equations. We demonstrate its potential applications in the context of general research questions in movement ecology using classical and new case studies as illustrations. We demonstrate the framework’s applications and versatility in classical home range models, but also show how it may be extended to address new ecological questions.

## Introduction

Space-use patterns stemming from animal movement behaviors provide signatures of organismal responses to ecological change. They contribute not only to our understanding of large-scale, population-level phenomena such as population and community dynamics, but also to policy development in conservation, natural resource management, and disease control (MacDonald and Rushton 2003; Hays et al. 2014; Epstein et al. 2020; Tao et al. 2021). Recent proliferation of high-resolution movement data enabled by advanced tracking technology has painted an increasingly dynamic picture of animal space-use patterns. Movement descriptions have consequently shifted from static summaries (e.g., what is the shape and size of an individual’s home range as defined by its long-term, stationary space-use pattern) to transient observations (e.g., how does an individual’s space-use pattern vary before it eventually stabilizes). This tendency toward explicit analyses of transient movement dynamics arguably represents a paradigm shift in movement research at large. Their feature of time scale as a central qualifier of space-use patterns assists in capturing animal movement responses timelier and more accurately in the wake of accelerated environmental disturbances. While transient analysis gradually becomes standard practice in empirical movement studies, spatially explicit movement models are slow to follow due to longstanding technical challenges in solving transient space-use dynamics. This paper introduces a numerical framework that that enables transient analysis of mechanistic movement models. We demonstrate its potential applications in the context of general research questions in movement ecology using classical and new case studies as illustrations.

Space-use pattern is commonly represented by utilization distribution (UD), which describes an animal’s probability of being at a location at a particular time. When studying area-restricted movement phenomena such as home range or territory formation, UD often refers to the equilibrium space-use pattern that the target animal presumably reaches and maintains during the surveillance period.

In theory development, individual-based models (IBM) have been one of the earliest approaches to address transient dynamics in UD, typically by simulating the evolution of movement trajectories of one or multiple individuals using correlated, biased random walks on a finite landscape. The step length and direction of each movement decision may be influenced by newly acquired environmental information as well as regular updates to the animal’s cognitive map of the environment (Van Moorter 2009; Riotte-Lambert 2015). However, since a spatially explicit pattern of usage such as a UD represents the full probability distribution of infinite stochastic simulations, each producing a single realization of movement trajectory, a sufficiently large number of random simulations must be performed to gain a clear, representative picture of how individual space-use evolves through time, which has so far been computationally prohibitive. Furthermore, they cannot directly evaluate how details in movement behaviors analytically affect temporal variations in space-use patterns.

Alternative IBMs have been developed to reconcile space-use variation with the underlying behavioral processes (Giuggioli et al. 2012; Potts et al. 2013; Potts and Schlägel 2020). The models are designed to uncover instability along territorial borders and characterize movement as random choices amongst nearest-neighbor lattices, excepting those sites that contain repellent environmental cues (e.g., scent-marks of conspecifics). The transient pattern of an animal’s UD, therefore, is equivalent to the changing distribution of its entries into all lattices when measured over a finite time window. Imposing specific movement rules reduces the model to a set of mean-field, partial differential equations that analytically relates specific behavioral descriptors (i.e., active scent time) to summary parameters of spatial dynamics (i.e., diffusion constant of territory border). The mathematics of this method is not yet fully understood, thus limiting the scope of its applications. Also, performing it is highly computationally intensive, making the models unwieldy for data fitting and testing ecological hypotheses.

Coincident with the developments of various IBMs has been the development of mechanistic space-use models. Mechanistic models seek to derive the time evolution of an animal’s UD from individual behavioral processes via the master equation, an iterative equation that describes the temporal relationship between successive UDs as a function of the animal’s responses to ecological feedbacks, including the preference to be near its cohort, avoidance of conspecifics, habitat selection, and departure from a previous cohort (Potts and Lewis 2014). The master equation may be subsequently converted into a time-derivative of UD expressed as a Fokker-Planck equation, wherein the dynamics of space-use patterns are separated into advection and diffusion components, akin to the collective properties of moving particles. This modeling approach has appeared in discrete and continuous forms in the seminal works of Okubo (1980) and Moorcroft and Lewis (2013). Solving the Fokker-Planck equation in time, however, is nontrivial whether through analytical or numerical methods. Therefore, by necessity, models have conventionally made the simplifying steady-state assumption by setting the time derivative to zero before solving for UD.

Steady-state analysis alone, however, may not give information about patterns of space-use over short ecological timescale, nor does it account for possible stochastic perturbations in the animal’s movement behavior. The recognition of this shortcoming, especially for its effect on the models’ predictive power, led to recent mechanistic models that approximate the transient dynamics of UD by solving for multiple steady states, each is driven by parameter values that represent the environmental condition at specific time (Moorcroft et al. 2006; Bateman et al. 2015; Tao et al. 2016). This *ad hoc* approach is still bounded by the steady-state assumption, but it also assumes the UD would stabilize faster than any behavior-altering changes in the environment, which implies a rate inequality that is often violated in reality (Benhamou 2014).

In this paper, we introduce to movement modelers a powerful software package that can derive transient UDs from a mechanistic model. Our easily reproducible template (see supplementary material) provides a practical guide to transient analysis at a comfortable level of abstraction where numerical algorithm is concerned. By pulling back the proverbial veil of steady-state solutions, we can glimpse new research possibilities that require direct detection of space-use patterns in flux, meanwhile helping to advance the burgeoning interest in transient dynamics amongst the broad community of movement ecologists.

We demonstrate the applications of transient space-use analysis by revisiting the classic model of scent-mediated conspecific avoidance, expanding on the original steady-state solution to give an unprecedented insight into the preceding transient dynamics of territory formation by each of the individuals. In later examples, we demonstrate transient UDs on a heterogeneous landscape, where structural barriers exist to alter space-use pattern during relocations. Finally, we apply transient space-use analysis to epidemiology and show how it may be used to test different intervention strategies amidst a growing outbreak.

## Methods

Mechanistic space-use models require descriptions of the underlying movement behavior (e.g., step length, turning angle) and scale-specific responses to environmental conditions (e.g., preference for resource-rich habitats, avoidance of foreign scent-marks). In animal home range and territory models, these mechanistic drivers are typically collected in the advection and diffusion terms in a Fokker-Planck equation:

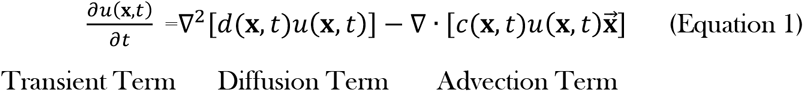

where *u*(**x**, *t*) denotes the individual’s UD at time *t*, 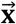 is a unit vector indicating the direction of the mean center of attraction relative to the individual’s current position. In the simplest case where a central point attractor is the only source of movement bias, *c*(**x**, *t*) and *d*(**x**, *t*) may be reduced to constant coefficients *C* and *d*. In that case, an exact steady-state solution exists (Moorcroft and Lewis 2013; Tao et al., 2016), assuming that the individual moves in radially symmetric manner within a finite 2-dimensional landscape Ω with zero-flux boundary condition.

Fokker-Planck is a special form of partial differential equation (PDE) where the solutions describe the time evolution of spatial random variables, such as the UD surface that characterizes individual spaceuse. FiPy (www.ctcms.nist.gov/fipy), developed for material science research by the National Institute of Standards and Technology (NIST), is a powerful Python-based PDE solver that we can use to solve for transients, i.e., transitional space-use patterns prior to the system reaches steady-state (Guyer et al. 2009).

Written in Python, FiPy inherits the language’s growing usage, easily legible syntax, and extensibility across its large libraries of algorithms for scientific computing (e.g., NumPy, SciPy) and visualization (Matplotlib). PDEs can be solved using FiPy function calls based on standard finite volume method, which reformulates the values of the dependent variable into discrete points on a meshed domain. The original PDE, thus reduced to a linear set of algebraic equations, can then be efficiently solved as a sparse matrix using an iterative scheme. The temporal solutions of the equation are contained within the cells that constitute the mesh.

### Background on Numerical PDE Solver

Several PDE solvers are available today, written for different language platforms and requiring varying levels of user experience with regard to numerical analysis. Given the diversity of options, however, few meet the specific criteria that make a tool suitable for an emergent group of interdisciplinary movement modelers. Several other Python-based solvers have recently appeared (Guyer et al. 2009), but difficult to operate for those researchers who prefer direct rendering of the governing equations without being distracted by lower levels of numerical detail. Many others (e.g., Matlab, Mathematica) are proprietary, costly, and lacking in versatility when it comes to visualizing the results. R has, in recent years, developed many powerful numerical methods (e.g., deSolve) that are syntactically familiar to ecologists and easy to implement at high level of mathematical abstraction (Soetaert et al. 2012). But for applications in space-use models, its PDE solvers suffer the major drawback of relying on the method-of-lines (MOL) instead of finite volume or finite element method. The procedure works by first discretizing only the spatial components of a PDE, thereby approximating it with a system of ordinary differential equations which are subsequently solved under discretized time. Although both approaches produce comparatively accurate solutions for 2-dimensional advection-diffusion problems (Selçuk et al. 2002), MOL’s inability to handle elliptic PDEs, i.e., when the time-derivative is removed, can preclude one from evaluating aspects of the transient dynamics with respect to the steady-state solution unless the latter is already known in its analytical form (but see R package ReacTran).

In contrast, FiPy comes ready with easy customization, relatively short set-up time to program, open-source nature, and capacity to solve large classes of PDEs. In addition, solution precision in FiPy can be further enhanced through a “sweeping” procedure. The overall runtime can also be shortened through parallel computing by exploiting the third-party package Trilinos. Studies of three-dimensional movement systems, commonly considered more appropriate for tracking avian and marine organisms, or those that require more topographically realistic spatial domain, can implement the simulations to run on complex mesh geometry coded separately in Gmsh.

Solving Fokker-Planck equation in time allows us to closely track the progression of an animal’s UD as it converges onto steady state. This dynamic approach broadens the type of movement problems we can mechanistically model, removing the traditional need to assume stabilized space-use. For instance, we may use Eq. 1 to model species introduction into an environment where the new home range attractor is arbitrarily located. Following the animals’ release, their subsequent space-use variation is observable via transient analysis, revealing significantly different resettlement processes depending on where the animals were initially distributed.

Transient space-use can be modeled in a landscape that is either homogenous or contains heterogeneous features that influence an individual’s local movement behavior, i.e., corridors connecting road-bisected habitats, or renewable resource that gets depleted over time. In our demonstrations of transient space-use dynamics, we assume movement is limited within a bounded landscape Ω with explicit edges. The gradient values along the perimeter are fixed at zero, thus ensuring that the area integrations of the UD solutions are maintained at unity.

We define Ω by constructing a mesh of either one- or two-dimensional space that consists of equidistant grid cells. At each time step, local UD solutions are solved at the cell centers, and the values between adjacent cells are estimated as flux across their boundaries (i.e., faces). When configuring the landscape, it is important to consider the tradeoff between spatial and temporal resolutions. According to the Courante-Friedriechs-Lewy (CFL) condition, increasing the number of grid cells shortens the maximal allowed time step, thereby increasing the computational time. However, placing point attractors at cell centers helps to keep the system dynamics robust to approximation error even under relatively low spatial resolution.

We optimize the precision of our UD solutions by considering different approximation schemes that are available for discretizing the advective term in the Fokker-Planck equation. In all our analyses, we adopt the explicit Van Leer flux splitting method, which achieves greater numerical precisions than several alternatives. For instance, hybrid, powerlaw, and related exponential-difference schemes, are more versatile to use but have been criticized for leading to qualitatively erroneous solutions under misalignment between the main advective direction and the mesh coordinates (Leonard and Drummond 1995).

## Results

### Extending classical case study: territory formation through scent-mediated conspecific avoidance

We perform transient analysis on well-studied mechanistic space-use models whose analytically derived steady-state solutions serve as the benchmark for assessing our time-dependent solutions. As a heuristic example, we revisit the scent-mediated conspecific avoidance model (White et al. 1995, 1996, Lewis et al. 1997, Moorcroft and Lewis 2006), where two territorial individuals, U and V, deposit scent-marks in mutual defense against foreign encroachment near their respective den sites at opposite ends of the landscape. Encounter with foreign scent marks causes each animal to respond by a) “over-marking” at a rate proportional to the local density of foreign scent marks, and b) increasing the likelihood of subsequent retreat toward its den site. This interaction between the UD and the scent-mark densities of both individuals can be modeled as:

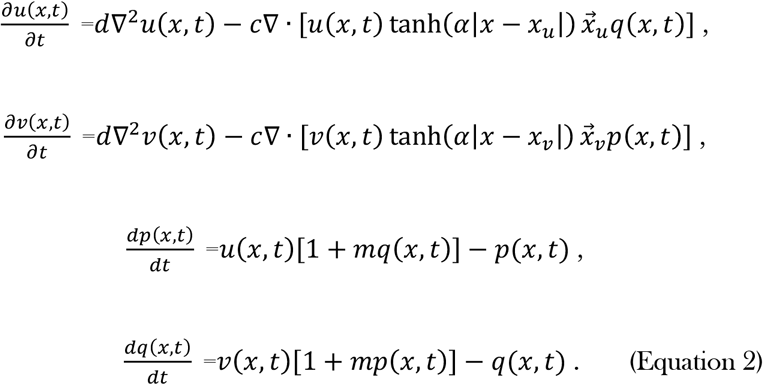

Here, *p*(*x, t*) and *q*(*x, t*) are nondimensionalized scent-mark densities of the two individuals, who are spatially distributed according to UDs, *u*(*x, t*) and *v*(*x, t*). 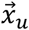 and 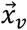 are unit vectors pointing toward their respective den sites. *m* denotes the strength of the overmarking response to foreign scent marks encountered. The inclusion of the hyperbolic tangent function removes the mathematical discontinuities at the point attractors to reduce numerical noise; *α* controls the smoothness of the step transition between opposite-signed unit vectors. We initialize both individuals using the same Gaussian UD centered on the origin and solve the transient dynamics of the system over a period of 10,000 time steps. Finally, we examine the system’s convergence toward previously published steady state solutions.

Our numerical solutions showed that the two scent-avoidant individuals released from the same site reproduced the territorial UDs at steady-states over time (Figure 1). The space-use patterns recreated two characteristic markers highlighted in the analytical work (White et al. 1996; Moorcroft and Lewis 2013): increased mutual avoidance of the inter-territorial boundary area, known as “buffer zones”, which is maintained by the abundance of scent-mark densities therein. The leveled parts of the individual UDs at two ends of the landscape indicate near-uniform movement directionality in areas where foreign scent-marks are absent. They are matched by values of the joint UD and the respective scent-mark densities.

**Figure 1.**
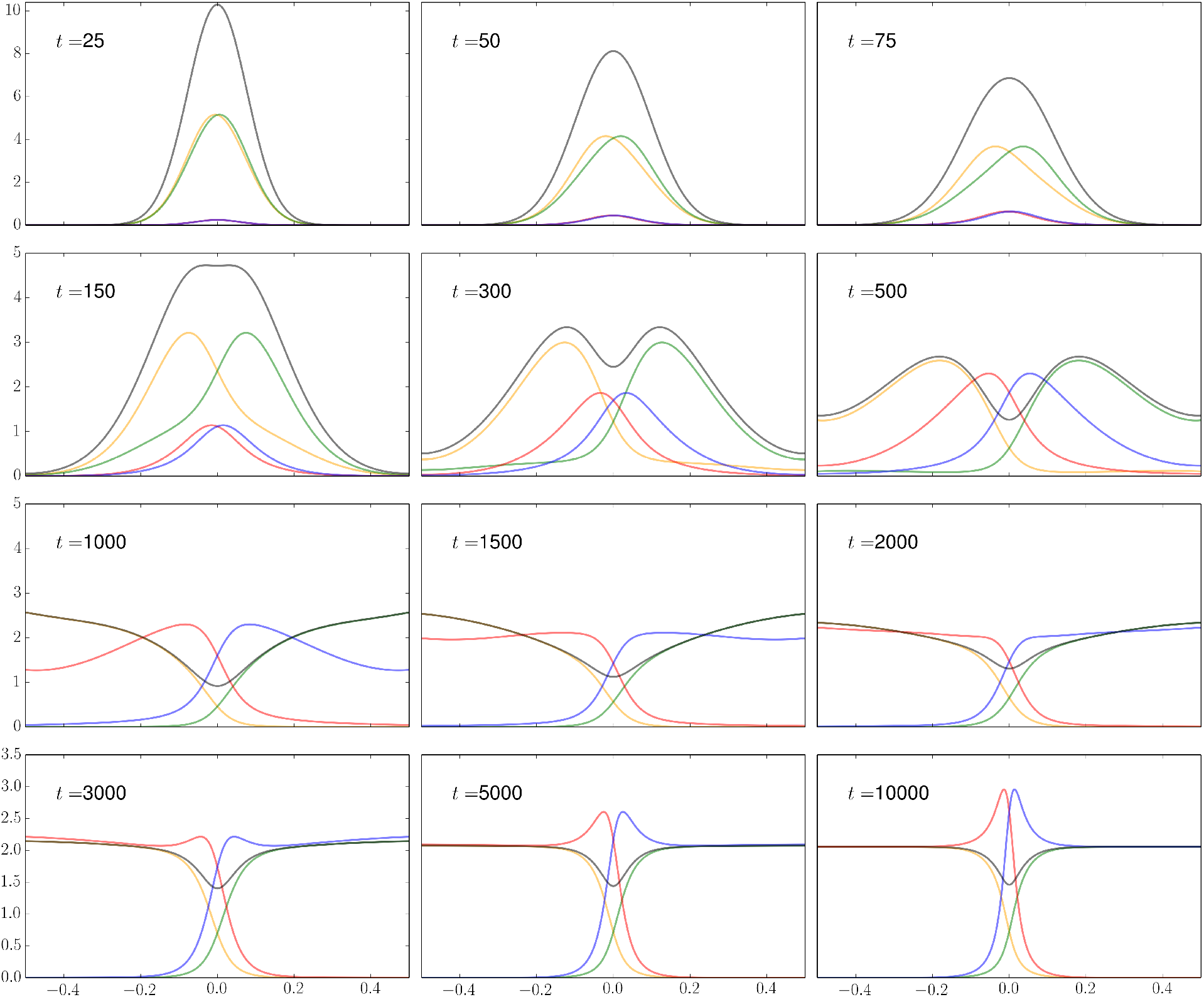
Territorial separation of two individual (or individual packs) on a one-dimensional landscape in avoidance of scent marks deposited by one another in the environment. Den sites are located at the two ends of the landscape. The yellow and green lines represent the individual utilization distributions; their sum is shown in black. The respective individual scent mark densities are displayed by the red and blue lines. The process of spatial segregation during the initial 10,000 time-steps post-release is captured by the transient dynamics of the utilization distributions; mutually deterrent boundary markings continue to accumulate long after the space-use convergence.

Transient space-use analysis revealed additional information about the space-use dynamics of the territorial neighbors. The visual time-series (Figure 1) showed that the buffer zones begin to appear relatively early during the process of neighbor separation (*t* = 150), concurrent with only a fractional buildup of the peak scent-mark density. Unexpectedly, while an animal is pushed toward its den site location, it briefly increases its presence in areas surrounding the conspecific den site, followed by its rapid removal as the owner’s scent mark residue accumulates (*t* = 300). We also observed that, for a period (*t* = 1,000), both individual and joint UD surfaces are highly skewed near the edges of the landscape before their shapes level to a plateau; meanwhile, the strength of the buffer zone (minimum of joint UD) starts to weaken until it settles to equilibrium (*t* = 2,000). Finally, once all the UDs have converged, the scent-mark densities in the buffer zones continue to increase for an extended duration (*t* = 5,000 to 10,000).

We can also extend the model to four territorial individuals interacting in a two-dimensional landscape, all initialized at the origin with a Gaussian UD and moving toward separate den sites under the same set of behavioral parameters. The large system of coupled equations (8 in total) is then solved in time for a period of 10,000 time steps. We follow the temporal variations in individual UDs and the accumulative scent mark densities for the whole group (Figure 2). At this higher spatial dimension and with more conspecific interactions, we uncovered a short-term ring-like pattern in the accumulative scent-mark distribution that quickly dissipated as the system approaches steady-state.

**Figure 2.**
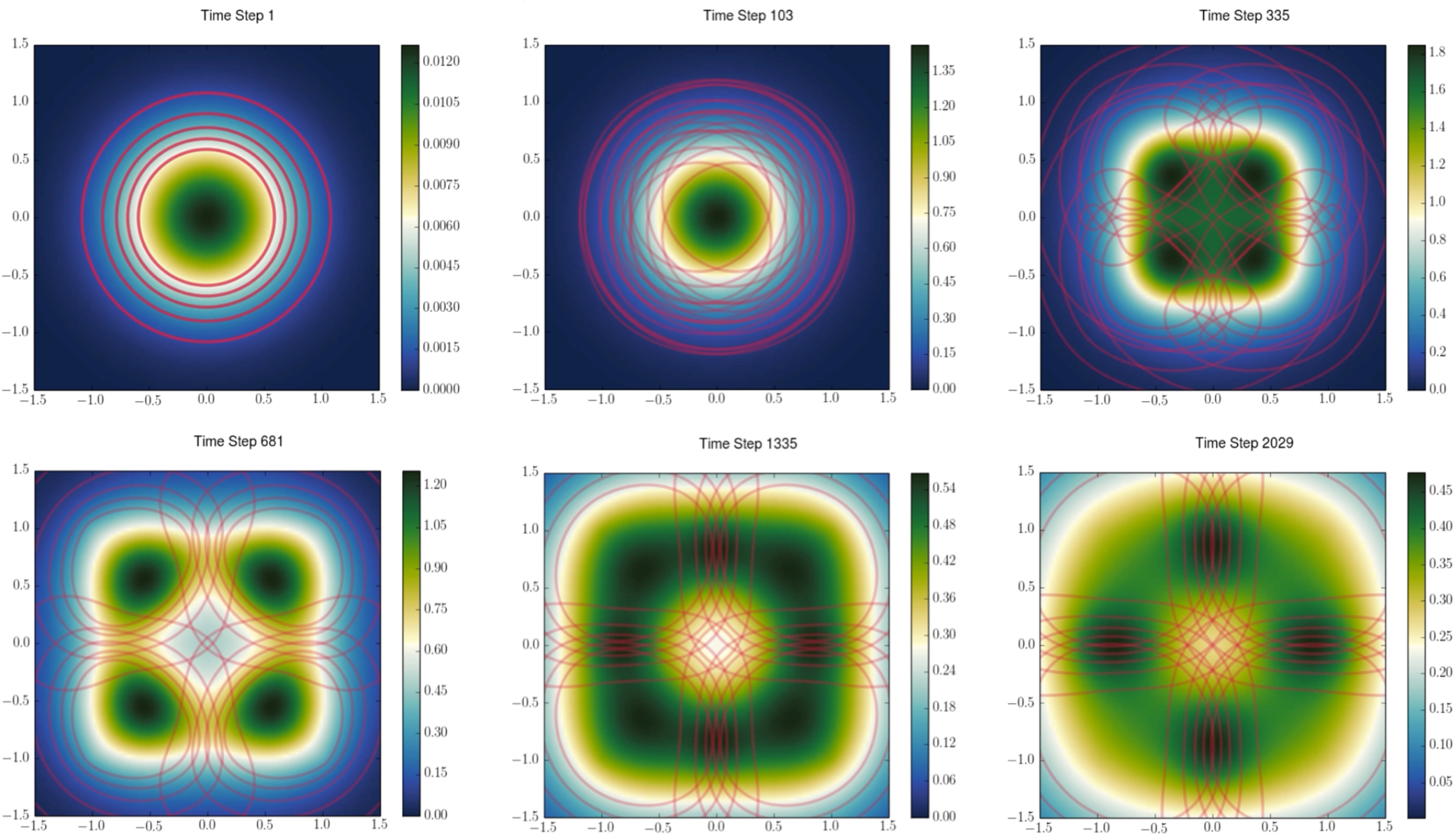
Territorial separation of four individual (or individual packs) on a two-dimensional landscape in avoidance of scent marks deposited by all others in the environment. Den sites are located at (±1, ±1). Red lines denote the contour maps of individual utilization distributions. The color density gradient shows the time-varying accumulative scent mark density.

### Incorporating heterogeneous landscape structures

The effects of spatial variation in landscape attributes (e.g., terrain steepness, forage density) on individual space-use patterns have been explored in classical movement models generally over long-term, i.e., in the form of steady-state UDs (Moorcroft et al. 2006; Bateman et al. 2015). However, omissions of short-term space-use patterns can undermine ecological inferences since it readily overlooks encounters with environmental hazards that can increase the risk of travel mortality (e.g., locally elevated rates of predation, starvation, and traffic collision). Using FiPy, we can closely monitor an individual’s whereabouts as it traverses across a heterogenous landscape and thus take immediate landscape effects into account.

To illustrate, we consider an animal returning to its shelter from its feeding ground. Passages are partially obstructed by a transecting roadway but open onto multiple movement corridors (i.e., overpasses) of equal width. We describe this system using the basic home range model in Eq. 1, initializing the UD as a Gaussian distribution at the feeding ground some distance away from the point-attractor. Now, however, the transient UDs are solved on a heterogenous domain divided by corridors into permeable and non-permeable regions. The landscape Ω is consequently constrained such that Ω = {**x**|**x** ∈ Ω_*T*_} with Ω_*T*_ indicating the contiguous traversable area, where zero-flux condition is enforced on both outer and inner boundaries *dΩ_T_*. The simple results (Figure 3) showed that while the UD eventually converges around the point-attractor as expected, for a period of time, the individual has nearly the same probability of being found at the shelter as in the middle corridors. Since time spent during transit correlates with an animal’s exposure to predation (Latham et al. 2011), this may help inform management decisions on where and how to build the movement corridors to maximize survivorship of the target individuals.

**Figure 3.**
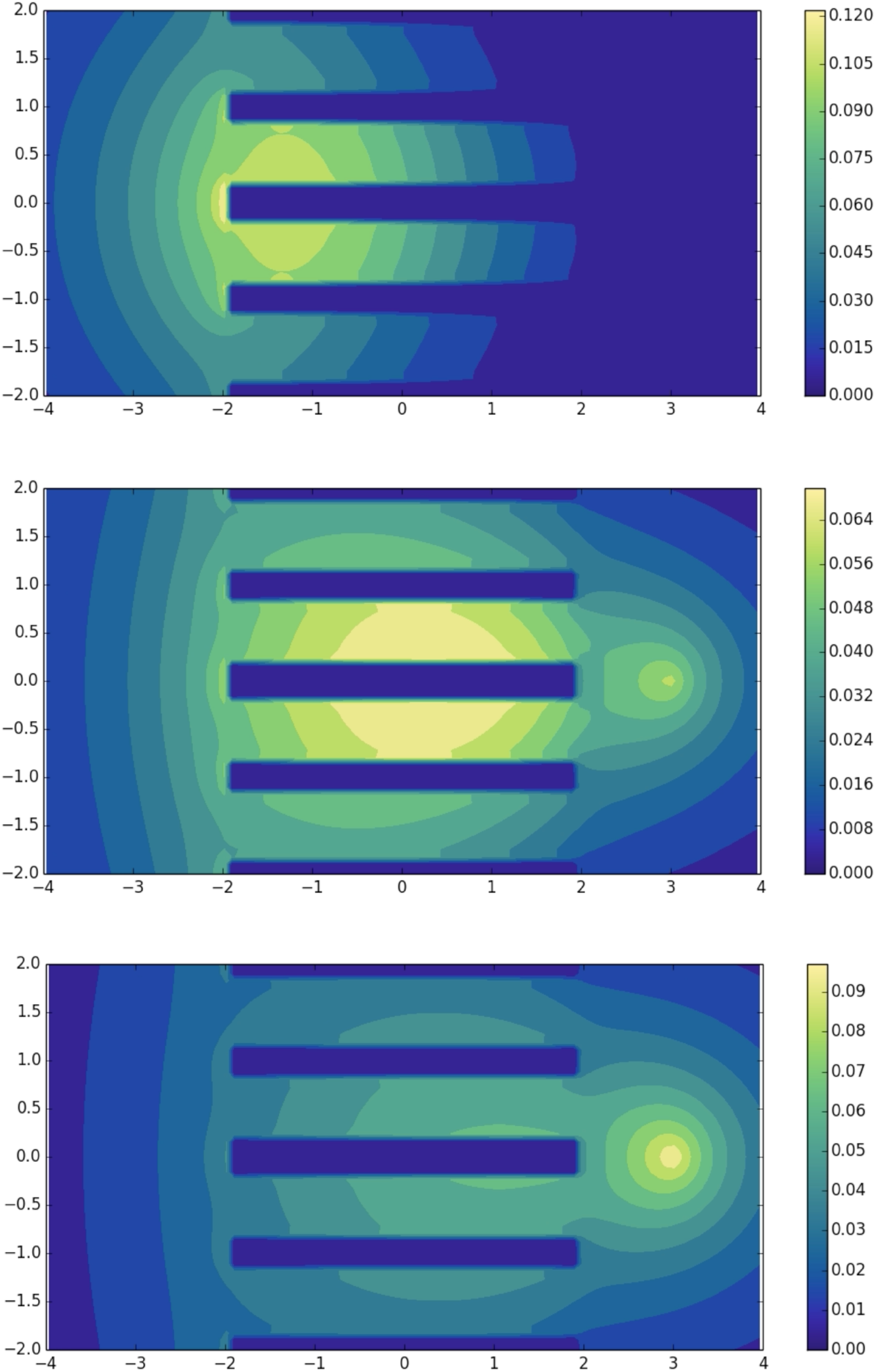
Transient space-use pattern of an individual relocating across a landscape connected by multiple parallel movement corridors. Its destination is centered around a point attractor located at the right-side of the landscape. The color density gradient shows the individual’s time-varying utilization distribution.

### Combining partial differential equations and individual-based models

Fokker-Planck equations alone are sufficient for predicting transient individual space-use dynamics in deterministic environments, where the spatial distributions of movement attractors, deterrents, corridors, and barriers remain constant or vary over time in a predictable manner. However, movement behaviors can also be induced or modulated by environmental noise, such as local occurrences of extreme weather events (Ziegler and Fagan 2014) and random outbreaks of infectious diseases inside the movement domain (Tao et al. 2021). We can directly address the effects of these stochastic processes on individual space-use patterns by developing a framework that couples and synchronizes the results of a spatial IBM and transient UDs.

To demonstrate this hybrid approach, we modeled vaccination rollouts as a public health response to a growing outbreak. Transmission and vaccination events are simulated on a two-dimensional landscape following a standard compartmental (susceptible-infected-vaccinated) model. Transmission occurs stochastically between infected and susceptible individuals with probabilities determined by a contact kernel. The rate that an individual receives vaccination declines with the local density of susceptible individuals on the basis that greater demand causes longer average wait-time. Vaccination personnel are deployed, i.e., moves, as a single unit governed by a basic Fokker-Planck equation. Its transient UDs, a representation of how control effort is distributed over space and time, are dependent on the deployment strategy that we can specify via the diffusion and advection terms. Here, the deployment strategy is guided by a mission to be present where vaccination coverage is maximized. We achieve this by reducing the diffusion coefficient with the local density of susceptibles, reflecting a strategic decision to expend more time on vaccinating densely populated neighborhoods. Furthermore, the advection coefficient (towards the initial deployment site) increases with the local density of infecteds, a mechanism we incorporated to enable redirections of vaccination effort away from neighborhoods that can no longer be protected.

Our results reveal that this particular deployment strategy can create a spatially asymmetric bubble of protection in which vaccination effort is concentrated and transmissions cannot readily occur (Figure 4). The temporal interplay between spatial disease dynamics and outbreak intervention can only be visualized by us concurring running the IBM and solving for the Fokker-Planck equation. Integrating these disparate modeling approaches allows us to predict the epidemiological consequences of implementing a robust set of deployment strategies amidst various realizations of a novel outbreak. Most importantly, this extension of our numerical technique enables practical, behaviorally complex management hypotheses to be tested, such as whether an outbreak can be better resisted by assembling the vaccinators into a defensive formation, or whether it is more effective to deploy healthcare teams into not-yet-infected areas beyond the epidemic front to impede spread.

**Figure 4.**
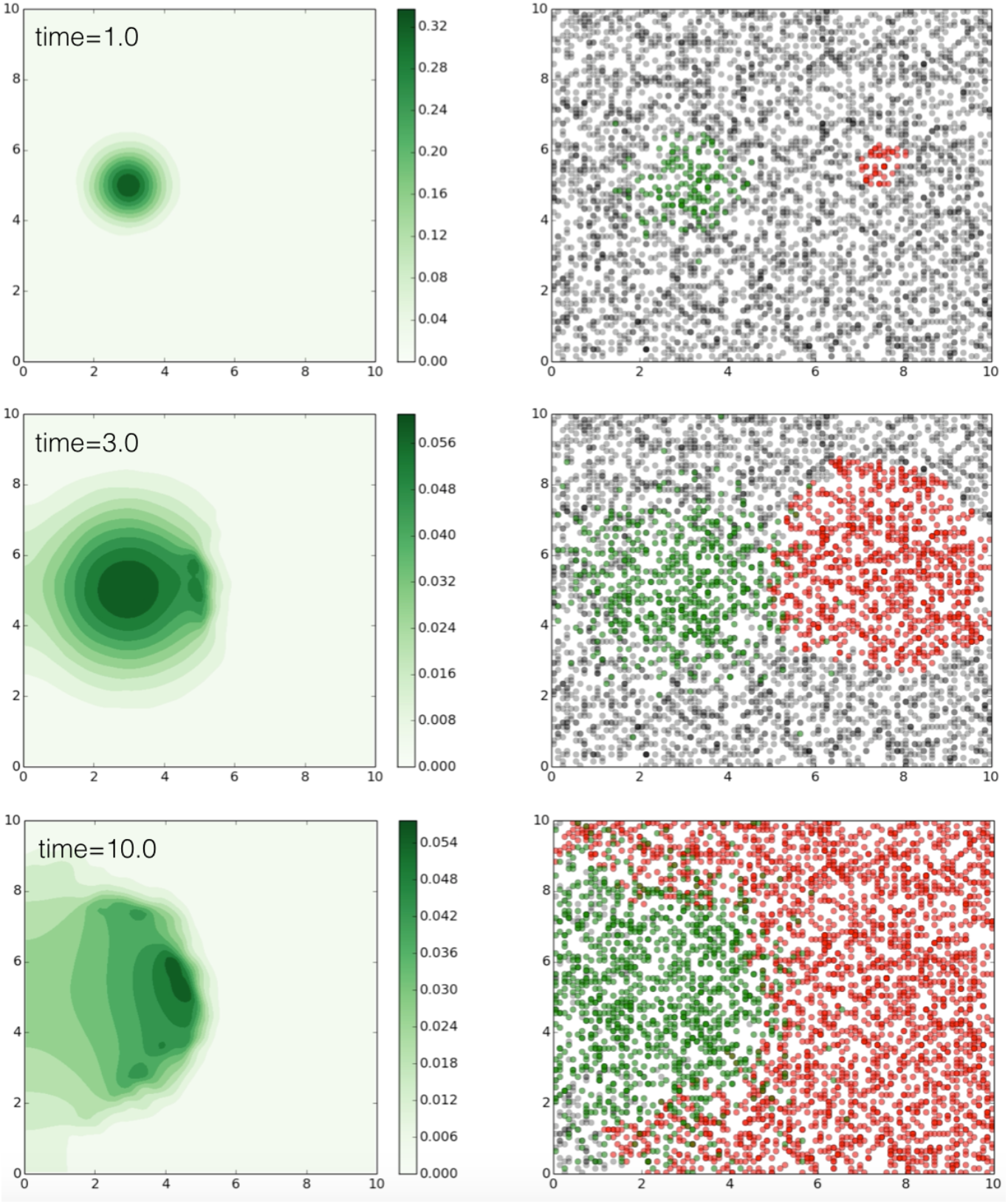
Disease control using a strategy based on behaviorally adaptive vaccine deployment, characterized by expending more control effort in high-needs neighborhoods and withdrawing from already infected regions. The green surfaces (left panels) depict the transient utilization distributions of the vaccination personnel in response to concurrent changes in local disease prevalence (right panels). Vaccinated, susceptible, and infected individuals are shown in green, grey, and red, respectively.

## Conclusion

In this paper, we aim to introduce a numerical modeling framework that can be used to solve for the elusive transient solutions of Fokker-Planck space-use models. The methods are relatively intuitive, self-contained, and can be implemented without extensive programming experience. We demonstrated the framework’s applications and versatility in classical home range models, but also showed how it may be extended to address new ecological questions.

Our example case studies suggest that transient movement dynamics deserve closer attention in the movement modeling literature. In systems where space-use pattern is slow to stabilize, transient utilization distributions may be the more accurate representation of an individual’s locality than the steady-states. Moreover, they can carry information that helps us gain a sequential view of pattern formation in movement and spatial ecology. Transient space-use analysis can therefore be used to draw more dynamical details from existing steady-state models while motivating new studies that focus on the ecological significance of transitional space-use patterns.

In this paper, we limited our examples to centralized movement governed by a single point attractor (e.g., den site, base of deployment). In natural systems, however, individuals might spend long periods of time moving between several coexisting point attractors. The modeling framework we have introduced can be further generalized to encompass such scenarios (e.g., multi-nucleic home ranges) in which the expected space-use pattern might be best described as a mixture of multiple transient utilization distributions driven by alternative directional preferences. An individual’s space-use pattern can also remain in a permanent state of transients due to increased sensory noise and long-term ecological disruptions, two immediate outcomes of global environmental change. Its ecological consequences remain an uncharted area of research.

Numerical solvers for partial differential equation could be a powerful and versatile tool for the next generation of mechanistic movement models. By bringing to light the transient utilization distributions from the models, they allow us to glimpse hidden dynamical features and challenge the conventional view of utilization distribution as a static measurement of movement.

## Supporting information

Practical Guide

