## Supplementary material for "Leveraging PDE solver for predicting transient space-use dynamics in ecological and epidemiological systems": Practical Guide

### Transient Mechanistic Home Range Analysis:

### A Practical Guide to Dynamical Modeling with FiPy Package

Yun Tao and Alan Hastings

Background

In today's "golden age" of movement ecology (Nathan et al. 2008), movement phenomena are currently gaining unprecedented ecological importance from a burgeoning interdisciplinary effort that aims to uncover mechanistic generalities across disparate systems, while generating reliable individual- and population-level predictions for specific organisms in response to vital management needs in the face of accelerating environmental change. Rapid development in tracking technology (Kie et al. 2010) has ushered in an influx of big movement data that can then be used to paint a more dynamically realistic portrait of movement patterns. Processed through advanced statistical methods, they promise to help us infer the temporal, or transient, effects of important movement determinants that have been previously overlooked when our observations were significantly coarse-grained. There is already growing evidence on the complexity of short-term animal movement dynamics, as illustrated by the interaction between individual state variable and its migration trajectory (e.g., Wittemyer et al. 2008), group cohesiveness over time (e.g., Pays et al. 2007), periodic fluctuation of territorial boundaries (e.g., Giuggioli et al. 2011), and seasonal variation in habitat selections (Avgar et al. 2013). It can be strongly argued that a conceptual paradigm shift towards movement transients is current underway (Tao et al. in prep).

Animal home range is becoming an active subject for transient analysis in the literature (e.g., Börger et al. 2006, 2008). Traditionally defined on the concept of an area-restricted space use that is invariant over extended periods of time, increasing availability of high-frequency sampling data is now challenging its former description whose quantitative basis relies on some extended timespans which are often subjectively assigned using rather coarse ecological delimitations, i.e., summer versus winter seasons (Kie et al. 2010). Home range estimates that fail to include a continuous time component are essentially omitting a level of pronounced dynamical detail that may be central to our understanding of the study system (Keeting & Cherry 2009).

Mechanistic home range models (Okubo 1980, Moorcroft and Lewis 2006), formulated from correlated random walks, have recently also started to address this conceptual shortcoming by incorporating temporal movement determinants (e.g., Bateman et al. 2014, Tao et al. 2014). Their Fokker-Planck framework models home range as a time-varying probability density function of an animal's location in space (i.e., utilization distribution), therefore allowing us to hypothetically track features of its space use pattern (e.g., topology, area coverage) as transient response variables to the system parameters related to demographic and environmental conditions.

These model, in their unsolved form, links the temporal variation in an individual's utilization distribution, u(x,t), to the directed (convective) and random (diffusive) motions with the following structure:

$$\underbrace{\frac{\partial u(x,t)}{\partial t}}_{\text{Transient Term}} = \underbrace{-\nabla \cdot [c(\mathbf{x},t)\, u(\mathbf{x},t)]}_{\text{Convection Term}} + \underbrace{\nabla^2 [D(\mathbf{x},t) u(\mathbf{x},t)]}_{\text{Diffusion Term}}. \quad (1)$$

The convection and diffusion coefficients, c(x,t) and D(x,t), describe specific movement rules that generally include the animal's mean travel velocity and directional bias towards its den site.

Although variants on this equation have been widely developed (Potts and Lewis 2014), their fundamental nonlinearity renders analytical solutions unfeasible and numerical estimates difficult to implement under many of the available computational tools, especially when more complex behavior and interactions amongst individuals are introduced. In contrast, the steady-state solution is much easier to obtain, but reflecting only the stable space use distribution when transient term reaches zero. This simplification created by removing the time derivative underlies the normative steady-state definition of home range and has to date remained largely uncontested, which is effectively prohibiting a differential analysis of home range transients. Rapid advancement in individual-based models (e.g., Wang and Grimm 2007) provides an alternative approach for generating temporal home range formations. However, these simulations, abound with complex movement patterns, lacks the burden of mathematical tractability. Furthermore, they are computationally limited in concluding any system-level generality that could be derived directly from the mean-field approximation implicit in Fokker-Planck equation. Tapping into the transient solutions locked within home range models would therefore deliver dynamical details along with a mechanistic understanding of their emergence, an effort that may in time facilitate an unification of the theoretical paradigms for animal movement analysis (Potts et al. 2014).

This paper introduces a powerful software package that is capable of deriving time-varying utilization distribution based on analytical equations. Our annotated and easily reproducible template provides a practical guide for home range researchers who wish to work above a level of abstraction where numerical algorithm is concerned. Our primary objective is to foster an emergent interest amidst the larger discipline of movement ecology and help raise questions related to the patterns and consequences of transient dynamics, and how the newfound access beyond the wall of steady-state can be used to advance our understanding of the complex interplay between mobile individuals and their environment. We will examine the potential for gaining conceptual insights from transient analysis solely based on the dynamical characteristics stemming from our heuristic examples. Lastly, we will demonstrate the software's application in classic territorial model of scent-marking conspecifics (White et al. 1996, Lewis et al. 1997) for which only the steady-state solutions have so far been determined.

<u>A Workflow of FiPy Home Range Simulation</u>

Fokker-Planck is a special form of partial differential equation (PDE) in which the solutions describe the time evolution of spatial random variables, such as the probability density surface that characterizes individual space use. FiPy (www.ctcms.nist.gov/fipy), originally developed by the National Institute of Standards and Technology (NIST) as a PDE solver for material science, is an open-source, Python-based software that we can leverage to remove the technical obstacle that has so far impeded transient analysis in studies of animal home range. Its choice of Python as the scripting language is motivated by latter's power, growing usage, easily legible syntax, and extensibility across its large libraries of algorithms for scientific computing (e.g., Numpy, Scipy) and visualization (Matplotlib). FiPy syntax complements Python's and provides straightforward function calls for customizing the PDE solver, which uses finite volume method that works by discretizing the space and time derivatives, thus casting a general PDE into a process that evolves incrementally over a mesh composed of contiguous polyhedral cells divided by facial boundaries. The original PDE, reduced to a linear set of algebraic equations, can then be efficiently solved as a sparse matrix using one of the built-in iterative schemes. The values of the dependent variable are contained within the cells, hence together forms a layer above the configured mesh. Further theoretical background on the algorithm can be found within FiPy's documentation.

Several PDE solvers are available today, written for different language platforms and requiring varying levels of user experience with regard of numerical analysis. Given the diversity of options, however, few meet the specific criteria that make a tool suitable for an emergent group of interdisciplinary movement modelers. Several other Python-based solvers have recently appeared (Guyer et al. 2009), but difficult to operate for those researchers who prefer direct rendering of the governing equations without being distracted by lower levels of numerical detail. Many others (e.g., Matlab, Mathematica) are proprietary, costly, and lacking in versatility when it comes to visualizing the results. R has in recent years developed many powerful numerical methods (e.g., deSolve, rootSolve) that are syntactically familiar to ecologists and easy to implement at high level of mathematical abstraction (Soetaert et al. 2012). But for applications in home range equation, its PDE solvers suffer the major drawback of relying on, instead of finite volume or finite element method, a technique called the method of lines (MOL). The procedure works by first discretizing only the spatial components of a PDE, thereby approximating it with a system of ordinary differential equations which are subsequently solved under discretized time. Although the accuracy of its transient solutions is comparable to the alternative approach for 2-dimensional convection-diffusion problems (Selçuk et al. 2002), MOL's inability to handle elliptic PDEs, i.e., when the time-derivative is removed, precludes one from evaluating aspects of the transient dynamics with respect to the steady-state solution unless the latter is already known in its analytical form. In contrast, FiPy comes ready with easy customization, relatively short set-up time to program, open-source nature, and capacity to solve large classes of PDEs. In addition, solution precision in FiPy can be further enhanced through a "sweeping" procedure. The over runtime can also be shortened through parallel computing by exploiting the third-party package Trilinos. Studies of three-dimensional movement systems, commonly considered more appropriate for tracking avian and marine organisms, or those that require more topographically realistic spatial domain, can implement the simulations to run on complex mesh geometry coded separately in Gmsh. Feeding a burgeoning community of FiPy users, the package is actively maintained by its core developers who also actively provide tailored technical support. Bug patches and additional features can be requested through the package's mailing list (ww.ctcms.nist.gov/fipy/mail).

We will demonstrate FiPy usage in home range analysis by presenting a general scripting template for solving equation 1 in one- and two-dimensional landscapes. Its transient capability is then put to the test when applied to a classic species-interaction model of territorial formation whose functions comprise a pair-wise coupled system of Fokker-Planck and ordinary differential equations. We infer the ecological implications of home range transients of heuristic examples and identify dynamical traits that would benefit from further investigations. Our first glimpse behind steady-state solutions motivates us to participate in the active dialogue raised by empirical researchers about the continuing relevance of traditional home range definition, both in concept and theory.

Transient Home Range in 1-Dimension

We illustrate FiPy applications in movement analysis through a set of fully working example code that solves for a series of transient home range solutions to equation 2, for a period leading up to convergence on the steady-state solution. Parameter names of FiPy modules are emboldened for clarity. For tutorial purpose, we first consider a one-dimensional system in which a recently released individual shows a directional preference for a point attractor some distance away. We outline the script according to a standard FiPy rubric that is structured according to the headings below. The original problem, its spatial domain, initial values, and boundary conditions are first defined, followed by numerical preconditions that control for the solutions' visual presentation, stability, and conservation throughout the simulated timeline.

*Domain Configuration*

The spatial domain in our example is a one-dimensional mesh defined on a grid of finite length $L$, consisting of $n$ number of uniform cells, whose size $dx = L/n$ equates the spatial resolution of our system and, for reason that will become apparent, affects the speed of the simulation. For a landscape symmetric about the origin:

```
>>> Mesh = Grid1D(nx=n, dx=L/n) + [[-L/2.]]
```

The term to the right of the plus sign permits our domain to extend halfway into the negative region.

Mesh construction determines the spatial arrangement of our solution. Each of its cells holds the dependent variable at its center, and the intermediate values between adjacent cells are approximated as flux across their partitioned boundaries, also known as faces. We choose to split our mesh along the x-axis into an odd number of cells such that the origin becomes positioned at a cell center. For L=3:

```
>>> Print Grid1D(nx=3, dx=3./3)+[[-1.5]].getCellCenters()[0]
[-1., 0., 1.]
```

Conversely,

```
>>> Print Grid1D(nx=4, dx=3./4)+[[-1.5]].getCellCenters()[0]
[-1.125., -0.375, 0.375, 1.125]
```

The appropriate number of cells is thus informed by the placement of the attractor, assumed generally to be at x=0. Its overlap with a cell center helps to ameliorate the effect of inadequate spatial resolution that would otherwise retrieve the critical point solution from poor face value approximation.

*Initialization*

Prior to start time of the simulated movement, our animal exists in a probabilistic space in the meshed domain. Its value at time zero is a function solely of spatial variable x, which we can access by setting

```
>>> x = Mesh.getCellCenters()[0].
```

We assume the individual have been moving randomly before exhibiting any home ranging behavior, and therefore initialize it using a Gaussian function centered at x=3 with standard variance, such that:

```
>>> init = 1/numerix.sqrt(2*numerix.pi) * numerix.exp((-((x-3.)**2/2)
```

Note that the use of numerix module, imported from FiPy, instead of numpy serves to avoid any conflict in namespace.

The utilization distribution u(x,t) can thus be defined as a cell variable u:

```
>>> u = CellVariable(mesh=Mesh, value=init)
```

*Fixing Boundary Condition*

Home range model is mathematically defined on an infinitely large landscape with zero-flux, reflective boundary condition along all its edges, thus keeping the integration of u(x,t) over all locations to unity throughout the movement process. Our numerical solutions are necessarily performed on a finite domain whose size relative to the solution should be sufficiently large to still meet the conservational condition. We correspondingly constrain the flux values on the left- and right-outermost faces bounding our 1-dimensional mesh by defining:

```
>> BCs = (FixedFlux(faces=Mesh.getFacesRight(), value=0.),
          FixedFlux(faces=Mesh.getFacesLeft(), value=0.))
```

*Transcribing PDE*

We assume our animal gravitates towards an attractor, $x_u = 0$, from both directions at constant speed c irrespective to its own location. The convective term can then be expressed as $c\,\tanh(\alpha x)$, which approaches unit step-function $c\,\mathrm{sgn}(x - x_u)$ as the smoothing parameter $\alpha \rightarrow$

∞. Because the convection coefficient indicating directed motion takes vector as argument, we assign a new spatial variable s valued as the facial flux of rank 1 over our mesh:

```
s = FaceVariable(mesh=Mesh, value=Mesh.getFaceCenters(), rank=1).
```

Equation 1 can now be transcribed into FiPy syntax, with parameters showing a convection-dominant process.

```
>>> alpha = 40
>>> c = 0.8
>>> D = 0.4
>>> eq = TransientTerm() == DiffusionTerm(coeff=D) +
                                        convection(coeff=(c*numerix.tanh(alpha*s)))
```

Note that to recognize the convection coefficient as a vector, it must take the form of a Python tuple as indicated by the extra parentheses. The convection function is currently a placeholder for specific scheme.

*Convection Scheme Selection*

It is worth clarifying, for the benefit of a general ecological readership, the differences between some of the common tools available in FiPy for discretizing the convection term, given that an inappropriate choice can severely undermine the solution's accuracy. For a time-dependent equation with high spatial resolution, we encourage using one of the several bundled explicit convection schemes (e.g., ExplicitUpwind, VanLeer) that achieve numerical precision albeit at the cost of longer runtime. We may sometimes also wish to perform a separate computation of the steady-state solution, by replacing TransientTerm with 0, in order to establish an independent reference alongside the transient dynamics when evaluating, for examples, a rate of convergence. In this case, explicit scheme becomes nonsensical and an implicit one must be applied. The ratio of convection to diffusion rates, defined as the Peclet number, informs the choice of implicit convection scheme. Hybrid, Exponential, and Powerlaw interpolate between extreme scenarios thus are suitable for most problems. For our purpose, we call on the second ordered Van Leer (explicit) scheme above the last line:

```
>>> convection = VanLeerConvectionTerm
```

*Stability Condition*

Due to the necessity of solving a continuous space, continuous time PDE over discrete spatial grids and time steps, scale resolutions in the space-time domains determine the solution's accuracy and stability. The Courante-Friedriechs-Lewy (CFL) condition explicitly prescribes the maximal length of time step that a solver tolerates when given the grid distance and convection coefficient. In one-dimensional system, we apply CFL by setting $dt = C\left(\frac{dx}{c}\right)$, with a value of $C$, known as the Courant number, equal to or less than 1 when we use an explicit time integration scheme typical for transient analysis.

We solve the original Fokker-Planck equation forwardly in time by an interval:

```
>>> dt = 1 * dx / c
```

*Plot Configuration*

For 1D and 2D visual display, FiPy uses Matplotlib to plot its the numerical results. We evoke the viewer function and passing over the dependent variable:

```
>>> viewer = Viewer(vars=u, limits={'xmin': -5., 'xmax': 6., 'ymin': 0., 'ymax': 1.3}
```

The figure window is framed to the dimensions set by the preferred axial limits.

*Chasing Transients*

We simulate a home ranging event over fixed duration in which an animal is orientationally biased towards the origin. To solve and observe the transient solutions in sequence, we set

```
>>> duration = 2000
>>> for step in range(duration):
>>>     eq.solve(var=u, boundaryConditions=BCs, dt=dt)
        viewer.plot()
```

The solutions can be plotted at each time step and their images stitched into a video format that directly illustrates the dynamics. Figures 1 and 2 show the 1D realizations of this model with two forms of initial distribution. In each, the transient home range, starting away from the den site, converges smoothly on the analytical solution (Moorcroft and Lewis 2006). We monitor the solutions' conservation to unity over time by outputting the product of mesh area and mean solution in each cell using the command:

```
>>> print u.getCellVolumeAverage() * Mesh.getCellVolumes().sum()
```

Transient Home Range in 2-Dimension

Extending the simulation to a more realistic description of terrestrial organisms, we can implement the same transient home range movement in a higher dimensional space after appropriate changes to the boundary and stability conditions explicitly enforced in the previous script.

For a two-dimensional landscape center around the origin, the mesh is created with a 2D grid function that allows the subsequent assignments of spatial variables:

```
>>> Mesh2D = Grid2D(nx=nx, ny=ny, dx=dx, dy=dy) + [[-l/2.]]
>>> x, y = Mesh2D.getCellCenters()
```

We can analogously initialize the cell variable u with a bivariate Gaussian distribution. The dimensional increase results in the inclusion of two additional mesh faces (top and bottom) in the fixed valued boundary condition, each of which should be combined into its neighboring face:

```
>>> facesTopRight = ((Mesh2D.getFacesRight()) | (Mesh2D.getFacesTop()))
>>> facesBottomLeft = ((Mesh2D.getFacesLeft())) | (Mesh2D.getFacesBottom()))
>>> BCs = (FixedValue(faces=facesTopRight, value=0.),
                (FixedValue(faces=facesBottomLeft, value=0.))
```

The convection coefficient now holds in nested tuple form the constant vector values in both x and y directions. Assuming isotropic movement:

```
>>> c = 1.0
>>> b = ((c,), (c,))
```

The CFL condition is likewise adjusted for two space dimension, under which

```
>>> dt = dx * 1. / (2*c)
```

The functional approximation of the convective flux term, under isotropy, acquires the form (White et al. 1995) $c = c_u \tanh(\alpha r)\frac{\mathbf{x}}{r}$, where $c_u$ is the convection speed and $r$ is the radius to the origin; $\mathbf{x}$ is represented by facial variable s as earlier. To perform arithmetic operation between r and $\mathbf{x}$, both are required to have the same array size, which we achieve by defining r also on facial positions. The Fokker-Planck equation is then transcribed as:

```
>>> xf, yf = Mesh2D.getFaceCenters()
>>> r = numerix.sqrt(xf**2+yf**2)
>>> eq = TransientTerm() == DiffusionTerm(coeff=D) +
    convection(coeff=(b*numerix.tanh(alpha*r)*s/r))
```

Snapshots of the transient movement towards the new home range are illustrated in figure 3.

Results

We illustrate default FiPy's capability in obtaining the transient solution to a general model of animal home range while heuristically keeping only the basic mechanism. Its simplicity notwithstanding, access to the numerical transients of the above example raises new research questions that potentially challenge some of the classic notions in movement ecology.

When we measure the area of home range given by the majority coverage of its space use distribution, we find that the sudden relocation towards distant attractor is accompanied by an initial diffusion in space use, which then immediately condenses into the time-independent distribution. To put it differently, the road to steady-state solution is spatially oscillatory. The expansion-collapse pattern captured only by transient solutions can easily apply to animals with rapidly moving home range center, which can be effected via foraging hotspots that periodically disappear and then reappear at remote locations, or movement response that lags notably behind the update rate of the animal's cognitive map that contains the locations of more favorable habitats.

Non-static home range center has been a developing component in more recent models that incorporate resource-selection. In populations that live under the perturbation of urban development, deforestation, resource removal, and habitat fragmentation, the inclusion of drifting attractors has critical applications in ecological management. But with little exception, the transitional space use pattern in the models is described by the steady-state solutions. Could this simplification where the transient dynamics is ignored mislead our prediction of movement consequence? To demonstrate the caveat of relying only on steady-state solutions, we simulate the above model in 2-dimension over combinations of 2 parameters: the distance between the two attractors, and the strength of convection. We then compute the extents of spatial variability prior to home range stabilization, as measured by the log-transformed variance of home range size throughout the relocation event. Figure 4 shows that proximity of new center correlates negatively with the magnitude of transient dynamics. When the convection strength is low, home range inflates initially with a diffusion component that is weakly reined, resulting in more significantly different transient pattern. However, transient dynamics again becomes pronounced when convection is increased beyond a certain threshold, a result that can be intuitively deduced from the greater contraction the home range undergoes at the new attractor. In general, our results show that, for predictive analysis involving individual home range pattern over a period of time, a complete reliance on steady-state solutions potentially dumps out crucial movement information between sampled dates.

Steady-state solutions of equation (1) are usually applied over arbitrarily defined timescales without scrutiny of the parametric effect on its degree of convergence. We evaluate the transient timescale as a function of model parameters. By numerically tracking the progression of transient solutions of individual space use towards the analytical steady-state that has been derived in earlier papers (Moorcroft and Lewis 2006; Tao et al. 2014), we directly compute the dynamical effects of several model parameters in governing the time required for home range pattern to stabilize. Prior to simulation, we hypothesize lengthening transient timescale with weaker convection strength $c$, stronger diffusivity $D$, and greater distance $l$ to the new attractor. The spatial trajectory of an individual that is initially released under a Gaussian distribution is subsequently tracked as it relocates based on the mechanistic description of equation 1. Convergence is observed in the increasing proportion of overlap between the probability density functions of transients and that of the steady-state solution. As expected (figure 5), home range emerges slowly in systems where movement is highly random, its directional bias constrained, and the destination far away.

Unpacking Complex Movement Transients

We so far illustrated the applications of FiPy environment for home range analysis using the foundational Fokker-Planck equation in which the movement rules are described by minimal ecological detail. We now put our numerical design to the test and explore the transient dynamics of well-established home range models whose analytically derived steady-state solutions serve as the benchmark to which our time-dependent solutions should ultimately approach. For our case study, we recall the scent-mark territorial models of two conspecifics (White et al. 1996, Lewis et al. 1997, Moorcroft and Lewis 2006), distributed by *u(x,t)* and *v(x,t)*, each governed by two behavioral responses upon encountering its neighbor's scent marks, *q(x,t)* and *p(x,t)*, respectively: increase in marking rate of its own by *m*, and higher propensity to turn back towards its own den site. This species-interaction movement model has the form:

$$\frac{\partial u}{\partial t} = D\nabla^2 u - c\nabla \cdot [u\vec{x}_u q] \qquad (4a)$$

$$\frac{\partial v}{\partial t} = D\nabla^2 v - c\nabla \cdot [v\vec{x}_v p] \qquad (4b)$$

$$\frac{dp}{dt} = u(1 + mq) - p \qquad (4c)$$

$$\frac{dq}{dt} = v(1 + mp) - q \qquad (4d)$$

The coupled nature of the above model plus the additional source terms in the bottom ODE portion creates numerical challenges previous analyses failed to resolve. In FiPy, we begin by simply defining and initializing four separate cell variables over the same mesh.

After assigning equidistant locations for both point attractors, the coupled PDEs is transcribed separately before being combined:

```
>>> eq1u = (TransientTerm(var=phi1) == DiffusionTerm(coeff=D, var=phi1) +
convection(coeff=(c * numerix.tanh(alpha*(s-den1)) * psi2.faceValue), var=phi1))
```

```
>>> eq2u = (TransientTerm(var=phi2) == DiffusionTerm(coeff=D, var=phi2) + convection(coeff=(c * numerix.tanh(alpha*(s-den2)) * psi1.faceValue), var=phi2))
>>> eq1p = (TransientTerm(var=psi1) == ImplicitSourceTerm(coeff=(1+ m*psi2), var=phi1) – psi1)
>>> eq2p = (TransientTerm(var=psi2) == ImplicitSourceTerm(coeff=(1+ m*psi1), var=phi2) – psi2)
>>> eq = (eq1u & eq2u & eq1p & eq2p)
```

The transient territorial formations are shown in figure 6, together with the analytical steady-state solution in the final panel. Several potentially interesting features of this process have not been previously witnessed. Even as the territorial space use separation between neighbors becomes apparent, the wall of scent-mark residue at that region undergoes significant qualitative changes over time, evolving from highly peaked to leveled surfaces, and eventually to the stationary shape that was derived analytically. This transient result suggests the possibility for computing the age of a pair of territories solely based on the coupled dynamics in their collective scent mark densities.

We can also simulate the process in two-dimensional landscape by modifying the mesh geometry and implementing isotropic convection term in equations 4a-d (figure 7). The resulting cell-fission-like dynamics over the course of territorial formations illustrates temporal variation in the topographical symmetry of each conspecific's inner home range structure as exclusive space use becomes increasingly visible.

Soetaert, Karline, Jeff Cash, and Francesca Mazzia. *Solving differential equations in R*. Springer, 2012.

Tao, Yun, L. Borger, and A. Hastings. "Dynamical Animal Home Range Analysis via Modal Selection Function: An Optimality Approach." *submitted.*

Tao, Yun, L. Borger, M. A. Lewis, and A. Hastings. "Towards Transient Movement Ecology." *In prep.*

Wang, Magnus, and Volker Grimm. "Home range dynamics and population regulation: An individual-based model of the common shrew *Sorex araneus*." *Ecological Modelling* 205.3 (2007): 397-409.

White, K. A. J., M. A. Lewis, and J. D. Murray. "A model for wolf-pack territory formation and maintenance." *Journal of Theoretical Biology* 178.1 (1996): 29-43.

Wittemyer, George, et al. "Disentangling the effects of forage, social rank, and risk on movement autocorrelation of elephants using Fourier and wavelet analyses." *Proceedings of the National Academy of Sciences* 105.49 (2008): 19108-19113.

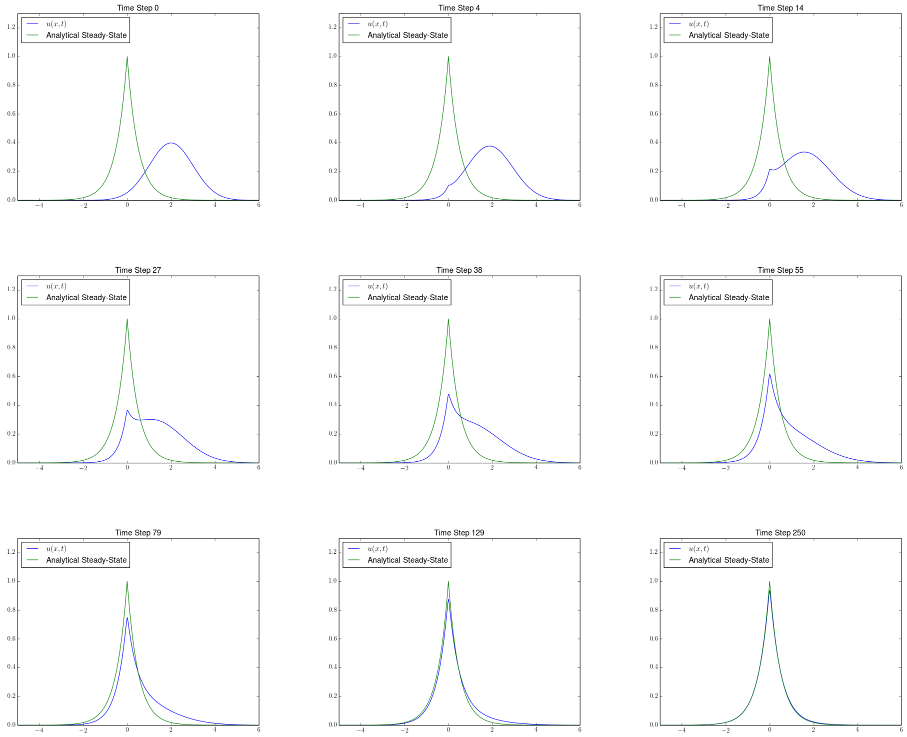


Figure 1: Snapshots capturing the time-evolution of home ranging space use towards a steady-state solution with a point attractor located at the origin (equation 3a). The individual is initialized as Gaussian distribution of $\sigma^2$=1 about $x = 2$. The mesh is a uniform 1D grid of length 24, consisting of 801 cells and fixed with zero-flux boundary condition. $D$, $c$, and $\alpha$ are 0.4, 0.8, 40, respectively. Van Leer convection scheme is chosen for the simulation over a time step that satisfies CFL with $C = 1$.

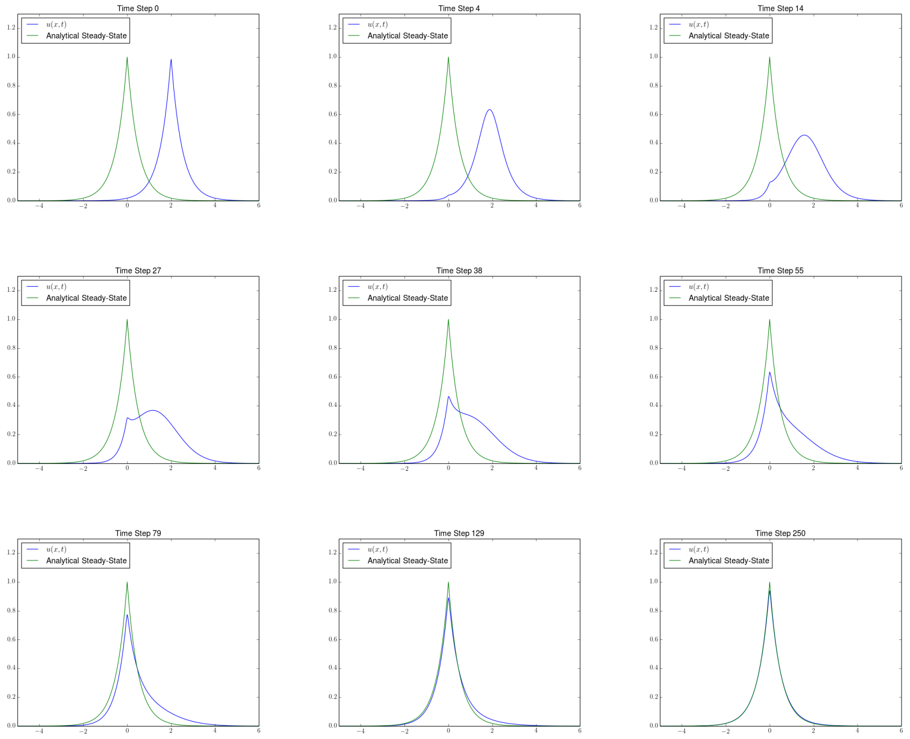

Figure 2: Analogous to figure 1 but with the initial condition set as a separate steady-state solution emergent from a point attractor located at $x = 2$ prior to the simulation. $D$ and $c$ are kept the same in the analytical functions of both steady-state solutions. All other parameters and configurations remain unchanged.

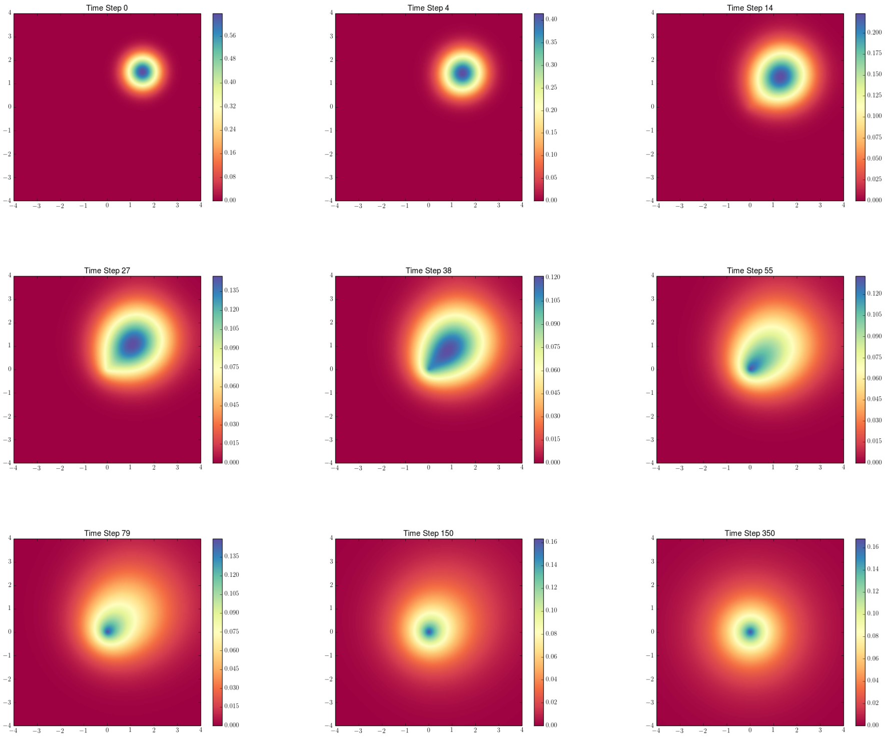


Figure 3: Snapshots of the transient space use of a home-ranging individual, who is initialized with a bivariate Gaussian distribution of mean position $(x, y) = (1.5, 1.5)$ and variance $\sigma_x^2 = \sigma_y^2 = 0.5$, moving towards its central attractor located at the origin. The mesh is a 2D grid with length 8 along each axis and made of 201x201 cells. Zero-flux boundary condition is implemented for all four edges of the domain. $D = c = 1.$, and $\alpha \approx 100$. Van Leer convection scheme is chosen for the simulation over a time step that satisfies CFL with $C = 0.2$.

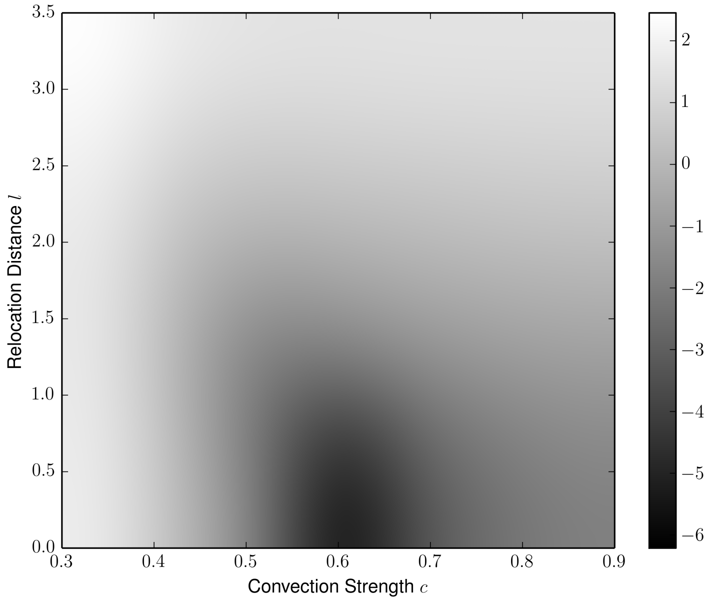


Figure 4: Log-transformed variance of home range size over time as a function of convection strength $c$ and distance $l$ between the animal's initial locational mean $(x, y) = (1.5, 1.5)$, with Gaussian variance $\sigma_x^2 = \sigma_y^2 = 0.5$, and its point attractor. Home range size is defined by the area under 90% utilization distribution. Movement occurs on a 101x101 cells, 2D grid of $L = 16$. $D$ is fixed at 0.2, and $\alpha \approx 100$. We simulate using Van Leer scheme and a CFL condition with $C = 0.3$ over 200 time steps.

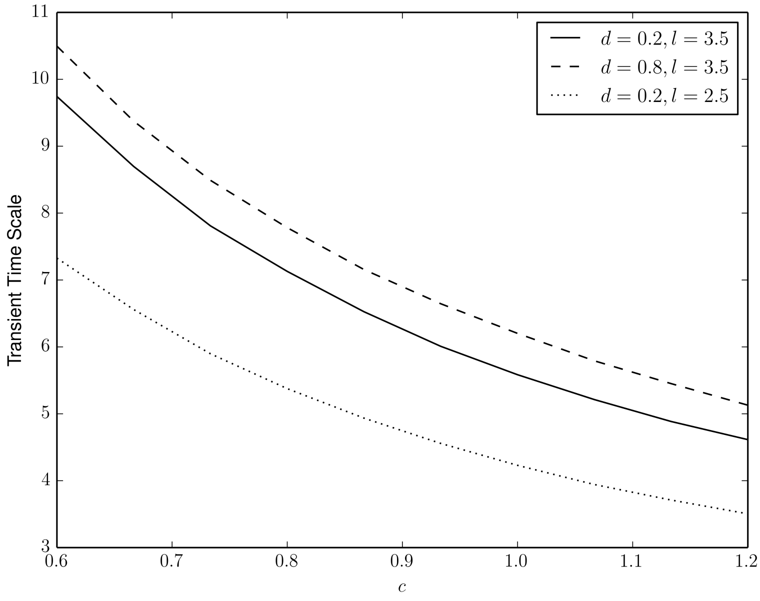


Figure 5: Standardized duration to 90% convergence of transients to steady-state spatial distribution in a 2D landscape as a function of model parameters. Mesh has length 16 and consists of 101x101 cells. The individual is initialized with a bivariate Gaussian distribution of mean $(x, y) = (1.5, 1.5)$ and $\sigma_x^2 = \sigma_y^2 = 0.5$. The attractor is located at the origin, with convection velocity set by $\alpha \approx 100$. Van Leer scheme is applied. 400 total time steps are simulated under CFL condition of $C = 0.3$. The functional form to the analytical solution is found in Tao et al. 2014.

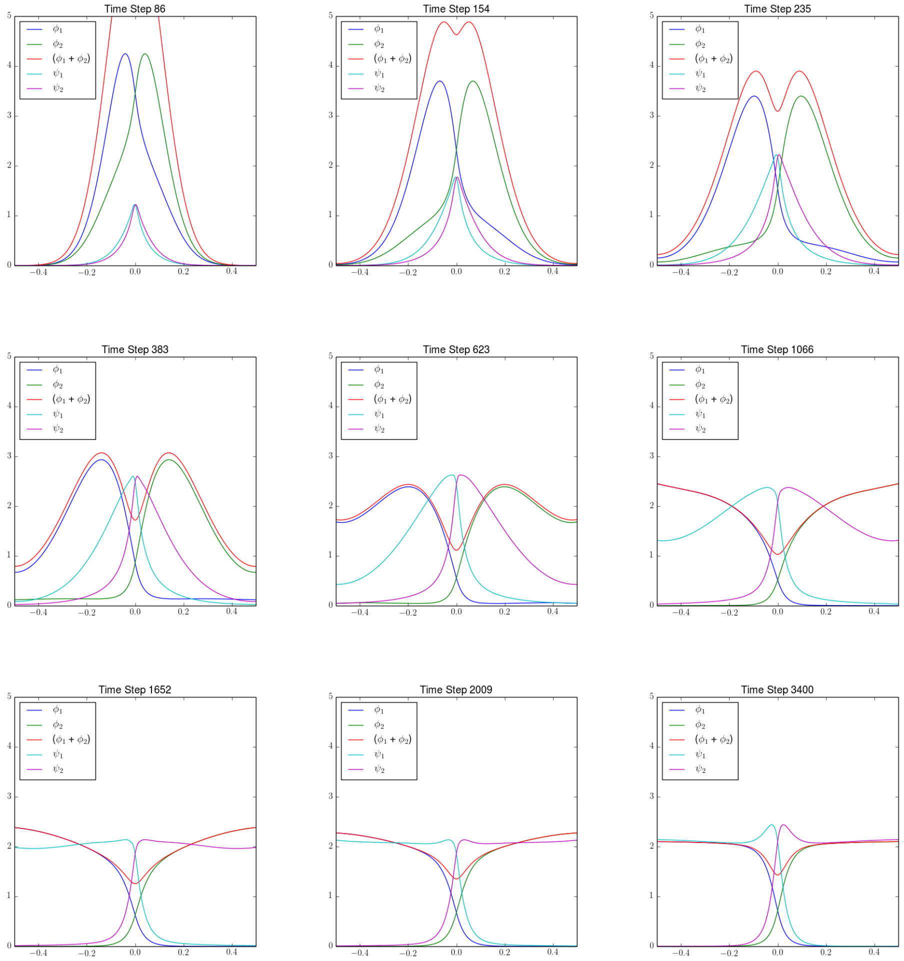


Figure 6: Transient 1D territorial formations of a pair of neighboring conspecifics, whose space use distributions $\phi_1$ and $\phi_2$ are initialized as a Dirac Delta function centered at the origin and their scent-mark density $(\psi_1, \psi_2)$ equaling zero. The mesh has length 1 consisting of 1001 cells. Den sites are equidistant from the origin, located at $x_1 = -1$ and $x_2 = 1$. Zero-flux boundary condition applies and conservation to unity is sustained. The process is convection dominant, with $D = 0.05$ and maximal convective speed $c = 0.7$ with notable dampening via $\alpha = 10$. Marking response parameter $m = 1$. Simulation runs on Van Leer scheme, satisfying CFL condition with $C = 1$.

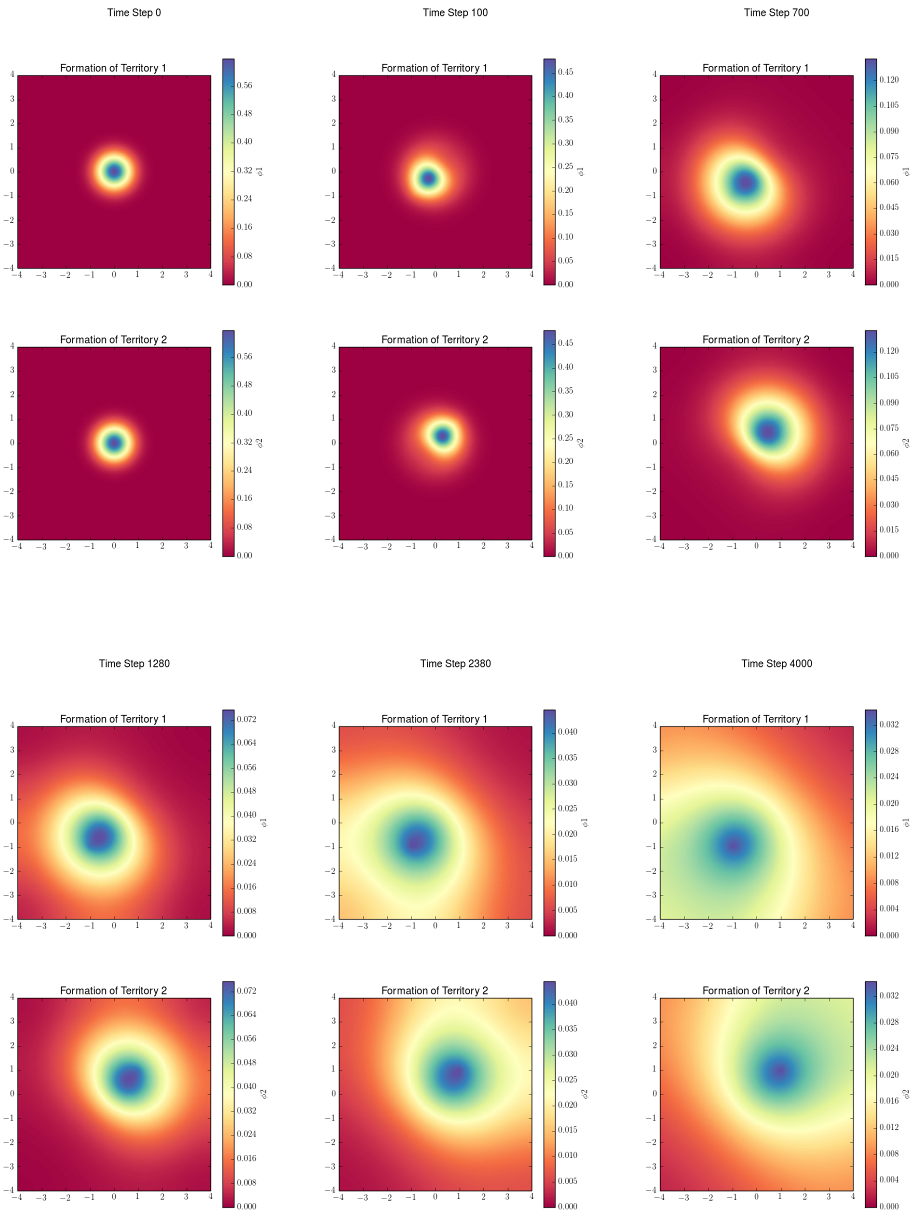


Figure 7: Transient territorial formations for a pair of neighboring conspecifics, both initialized with the same origin-centered bivariate Gaussian distribution of isotropic variance $\sigma_x^2 = \sigma_y^2 = 0.5$. Their respective den sites of radially symmetric convective forces are located at $(x_1, y_1) = (-1, -1)$ and $(x_2, y_2) = (1, 1)$. Scent-mark density is set to zero for both animals at start time. The mesh is a 2D grid with axial length $l = 8$ and cell size of 201x201. We configure zero-flux boundary condition and conservational constraint is maintained throughout simulation. $D_1 = 0.05$; convective speed $c = 1$ with $\alpha \approx 100$. Scent-mark parameter $m = 0.02$. Van Leer convection scheme is selected along with a time step that satisfies CFL with $C = 1$.
